# Development and validation of ultra performance liquid chromatography tandem mass spectrometry (UPLC-MS/MS) method to quantify monotropein in blueberries

**DOI:** 10.1101/2025.07.23.666277

**Authors:** Ishveen Kaur, Courtney P. Leisner, Dennis P. Cladis

## Abstract

Blueberries (*Vaccinium* species) are an economically important fruit crop rich in bioactive compounds like polyphenols and flavonoids. Interestingly, some blueberry cultivars also produce monotropein, which has bioactive properties, including anti-inflammatory and neuroprotective effects. However, methods to quantify monotropein in blueberries have not been optimized. To address this gap, an optimized analytical method for monotropein extraction and quantification using liquid chromatography tandem mass spectrometry (LC-MS/MS) was developed. Different extraction strategies were compared, including variations in temperature, time, and ultrasonication treatments. Optimal extraction was achieved by heating samples to 60°C for 15 minutes in methanol. The method had high percent recovery (89-110% intraday; 91-108% interday) and good repeatability (1.17-2.15% relative standard deviation (RSD) intraday; 4.68-7.16% RSD interday). This protocol was then applied to 28 blueberry cultivars, 14 of which had not been previously analyzed for monotropein. Monotropein ranged from 0-1807 ng/mg dry weight. The developed method provides a robust tool that can be applied to future evaluations of monotropein in diverse blueberry cultivars.

## 1. INTRODUCTION

Plants produce two types of metabolites, broadly categorized into primary and secondary (specialized) metabolites. Primary metabolites are integral to plant growth, development, and reproduction [1]. These compounds are directly involved in energy production and contribute to plant growth and yield [2, 3]. Plants also produce specialized metabolites that enable them to respond to environmental changes and can play key roles in defense against pests, pathogens, insects, and herbivores [4, 5]. Most of these compounds are synthesized from intermediates of primary metabolic pathways. These plant specialized metabolites are broadly characterized into three main categories based on their function, structure, and biosynthesis: terpenes, (poly)phenols, and alkaloids [6].

Terpenes, characterized by the combination of one or more isoprene (C_5_H_8_) units, are the largest class of specialized metabolites, with over 30,000 unique compounds identified [7]. Iridoids are a subclass of terpenes that have the general form of a cyclopentanopyran [8]. Iridoids are widely distributed across various plant families and are typically found in the glycosylated form *in planta* [9, 10]. Plants synthesize iridoids in response to biotic and abiotic stresses, protecting plants against herbivory, pathogen attacks, and environmental challenges such as temperature stress, extreme heat, temperature, UV, light, radiation [1]. When consumed by humans, iridoids may have health promoting properties, including anti-inflammatory, neuroprotective, antioxidant, antidiabetic, antimicrobial and cardioprotective activities [11, 12].

Monotropein is an iridoid glycoside recognized as the principal bioactive compound in *Morinda officinalis* (family Rubiaceae), a plant widely used in traditional Chinese medicine (TCM) for the treatment of osteoporosis and inflammation [13-15]. Traditionally extracted from the roots of *M. officinalis*, monotropein has been reported to exhibit a broad spectrum of pharmacological activities, including antioxidant, anti-inflammatory, and anti-apoptotic effects [16, 17]. Beyond TCM applications, monotropein is also employed in Polynesian ethnomedicine to manage a range of ailments such as infections, diabetes, asthma, hypertension, and pain [16]. These diverse therapeutic uses underscore the potential of monotropein as a multifunctional, medicinal, natural product.

Recent studies have identified the presence of monotropein in blueberries (*Vaccinium* species) [18]. Blueberries are an economically important fruit crop valued for their abundant phytochemical content and notable antioxidant activity [18-22]. In a comprehensive metabolite screen of an 84-member *Vaccinium* diversity panel, Leisner et al. [18] reported that monotropein was detectable in all 13 wild *Vaccinium* species analyzed but only 5 out of 71 cultivated blueberry accessions, suggesting that monotropein may be lost through cultivation of commercial blueberries [18].

While the Leisner et al. [18] study was the first to report the presence of monotropein in cultivated blueberries, the extraction protocol was not optimized. Therefore, the goal of the present study was to develop and validate a robust analytical protocol to quantify monotropein across different blueberry varieties. A systematic approach was used to optimize extraction efficiency, including the use of various temperatures, extraction durations, and the use of ultrasonication. After optimization, the protocol was applied to the quantification of monotropein in 28 blueberry accessions, many of which have never been analyzed for monotropein content.

## 2. MATERIALS AND METHODS

### 2.1 Materials and supplies

#### 2.1.1 Plant samples and materials

All methods were developed using fruit from a lyophilized, monotropein-positive wild blueberry composite (WBC) supplied by the Wild Blueberry Association of America (WBANA). The optimized protocol was then applied to *Vaccinium* fruits (both wild and cultivated) obtained from the United States Department of Agriculture Germplasm Resources Information Network (USDA GRIN), in Corvallis, Oregon, USA. The samples collected from USDA GRIN represent *Vaccinium* spp. from several different countries of origin; accession identification information is shown in Table S1. Samples were shipped from GRIN overnight on ice, flash frozen, and stored at -80°C until analysis. A single biological replicate was obtained for all *Vaccinium* spp. from USDA GRIN. Additionally, samples of cultivated blueberry were also analyzed. This material was obtained from various public nurseries and breeding programs.

#### 2.1.2 Chemical materials and supplies

A commercial standard of monotropein was purchased from Sigma-Aldrich (5945-50-6, St. Louis, MO, USA). LC-MS grade solvents, including methanol, acetonitrile, and formic acid were purchased from Thermo Fisher Scientific (Waltham, MA, USA). Type I ultrapure water (18-2 MΩ.cm) was used for all experiments (PureLab Flex, Elga Veolia, High Wycombe, United Kingdom).

### 2.2 Extraction optimization

For extraction optimization, 50 mg of WBC was mixed with 5 ml of LC-MS grade methanol and vortexed for 1 minute. After vortexing, the samples were subjected to different treatments as described below and shown in Table 1. Samples were then centrifuged (Beckman Coulter Allegra V-15R, Brea, California, USA) for 5 minutes at 1776x *g*; the supernatant was decanted and the remaining pellet extracted a second time. The supernatants were combined and dried completely under nitrogen using a nitrogen evaporator (Organomation N-Evap 116, Berlin, Massachusetts, USA). Dried extracts were resolubilized with 100 uL methanol and vortexed for 1 minute. After vortexing, 1900 ul of water was added before vortexing again. The resolubilized extract was filtered through 0.45 um PTFE filters into LC vials and analyzed via LC-MS/MS, as described below.

**Table 1.** Extraction parameters tested.

| Parameter | Variables tested |
| --- | --- |
| Time | 15min, 30min, 45min, 60 min |
| Temperature | Room Temperature, 30°C, 45°C, 60°C |
| Heat x sonication | Heat alone, Sonication alone, Sonication followed by Heat, Heat followed by Sonication |

Different parameters were used to optimize the extraction protocol for monotropein quantification. Heat, ultrasonication, and combination of both at different temperatures and time intervals were analyzed to optimize monotropein extraction (Table 1). For heating, a water bath (Fisherband-GPD05, Hampton, NH, USA) was used. Heating for different time intervals (15, 30, 45 and 60 minutes) and at different temperatures (room temperature, 30, 45, and 60°C) to extract monotropein from the blueberry fruit tissue were evaluated. Additionally, an ultrasonication method was used to disrupt the cells using high frequency ultrasonic waves emitted by a sonicator (Thermo Fisher Scientific, FS30D Ultrasonic Cleaner). Heat and ultrasonication methods were used independently as well as in different combinations to find the most effective method for a total of thirty-three treatment combinations for (see Table 1 for complete list).

### 2.3 UPLC-MS/MS analysis

Monotropein was quantified via Ultra-Performance Liquid Chromatography tandem Mass-spectrometry (UPLC-MS/MS) using a Waters UPLC Acquity H Class system equipped with a Xevo TQ-S micro detector (Waters Corporation, Milford MA, USA). Samples were injected and separated using an Acquity BEH C18 column (2.1 um, 1.7 mm id x 50 mm) with a flow rate of 0.5 mL/min. Samples were eluted using a biphasic gradient of solvent A (0.1% formic acid in water) and solvent B (0.1% formic acid in acetonitrile) as follows: 0 min, 0% B; 1.0 min, 5% B; 1.5 min, 95% B; 2.5 min, 95% B; 2.6 min, 0% B; 4.0 min, 0% B. MS conditions were as follows: capillary voltage, 0.5 kV; source temp, 150°C; desolvation temp, 600°C; desolvation gas flow, 1000 L/hr; cone gas flow, 50 L/hr. Identification and quantification of monotropein was based on the authentic standard, using a calibration curve ranging from 0.001-100 µg/mL. Monotropein (390.3 g/mol) had a retention time of 1.04 min and was detected using electrospray ionization operating in positive mode (ESI+) coupled with a triple quadrupole mass spectrometer. The parent ion (m/z 413.1) was detected as the sodium adduct of monotropein ([M+Na]^+^). A total of four mass transitions were optimized using IntelliStart software (413.117 → 185.020, 413.117 → 202.978, 413.117 → 233.036, and 413.117 → 251.056; Fig S1-S2). The cone voltage for all transitions was 26 V, and the collision energies were 22, 26, 26, and 22 eV, respectively. The most abundant fragment (413.117 → 233.036) was used for quantification (Fig S1-S2).

### 2.4 Method validation

#### 2.4.1 Instrument detection limit (IDL), method detection limit (MDL), and limit of quantification (LOQ)

Limits of detection and quantification on the LC-MS/MS were estimated based on the standard deviation (σ) of five replicates each of five concentrations of the monotropein standard (0.004, 0.01, 0.04, 0.1, and 0.4 μg/mL), where the IDL was estimated as 3σ and LOQ_LCMS_ was estimated as 10σ, as previously described [23, 24]. To back calculate the MDL and LOQ_blueberry_ to determine the detection and quantification limits in lyophilized blueberry fruits, the IDL and LOQ_LCMS_ were multiplied by the resolubilization volume (i.e., 2 mL) and divided by the mass of extracted blueberry tissue (i.e., 50 mg), as previously described [24].

#### 2.4.2 Repeatability

Repeatability was determined as previously reported. Briefly, intraday repeatability was assessed by analyzing 3 concentrations of the monotropein standard (0.1, 1, and 10 μg/mL), 5 times on a single day. Interday repeatability was determined across 5 consecutive days by analyzing 3 concentrations of the monotropein standard (0.1, 1, and 10 μg/mL) 3 times per day. Repeatability was determined as the relative standard deviation (%RSD): (standard deviation/mean) x 100%, as previously described [24-26].

#### 2.4.3 Percent recovery

Recovery was determined using three concentrations of the monotropein standard (0.1, 1, and 10 μg/mL) added to either blanks or WBC powder. Each was extracted in triplicate to determine recovery, as previously described [25, 26]. After extraction and analysis, the % recovery was calculated by: (measured amount of monotropein standard)/(starting amount of monotropein) x 100%.

### 2.5-Statistical analysis

Data were analyzed for outliers, normality, homogeneity of variance and collinearity by analyzing homogeneity of variance, goodness of fit, and generating Q-Q plots. Due to the non-normal distribution of the data, the Kruskal–Wallis (KW) test was used. KW is a non-parametric alternative to one-way ANOVA and was used to evaluate differences between extraction strategies. The generalized linear model included time (15, 30, 45, and 60min), treatment group (heat, sonication, heat then sonication, and sonication then heat), and temperature (room temperature, 30°C, 45°C, and 60°C) as variables. All statistical analyses were performed in R version 4.3.1 [27]. To determine differences between monotropein content in wild versus cultivated varieties, the non-parametric Mann-Whitney test was used to due to the non-normal distribution of the data.

## 3. RESULTS

### 3.1 Optimization of monotropein extraction conditions

To systematically optimize the extraction protocol, several techniques were used, including sonication, heat, and time (Table 1). These parameters were chosen based on previous extractions of specialized metabolites, including monotropein [18, 28]. These extraction parameters were then systematically evaluated to determine the most efficient conditions. To start, heat alone, sonication alone, and the combination of both were compared. There were no significant differences in extraction efficiency (*p* > 0.05). Therefore, to streamline the method, the most time- and labor-intensive options (i.e., the sequential combinations of heat and sonication, regardless of their order) were eliminated.

Then, the effect of extraction time (i.e., 15, 30, 45, or 60 min) for the heat-only and sonication-only treatments were examined. Extraction time had no significant effect on extraction efficiency for either technique (*p* > 0.05). Thus, for both heat and sonication, only the 15-minute treatment (i.e., the most efficient option that did not compromise extraction efficiency) was considered further. When evaluating the effect of temperature (i.e., 30°C, 45°C, or 60°C) on extraction yield for the 15 minute, heat-only treatment, the 60°C treatment demonstrated significantly higher monotropein extraction (*p* < 0.05), making it the most favorable condition. Finally, a direct comparison between the best heat-only condition (i.e., 60°C for 15 min) and the best sonication-only condition (i.e., 15 min) showed no significant difference (*p* > 0.05). Therefore, due to the simplicity, ease of application in most laboratory settings, and to maintain consistency with previous reports of monotropein extraction from blueberries [18], heat-only at 60°C for 15 minutes was selected as the optimized method.

### 3.2 Method validation

#### 3.2.1 Sensitivity and limits of detection and quantification

After optimizing the extraction method, we evaluated the sensitivity of the LC-MS/MS method to detect monotropein (Table 2). The instrument detection limit (IDL) is the smallest concentration of monotropein that can be detected on the LC-MS/MS, while the limit of quantification (LOQ_LCMS_) is the smallest concentration that can be quantified. Similarly, the method detection limit (MDL) is the smallest amount of monotropein that can be detected in the lyophilized blueberries themselves, while the limit of quantification (LOQ_blueberry_) is the smallest concentration that can be quantified. The IDL and LOQ_LCMS_ were 0.0103 and 0.0273 μg/mL, respectively, while the MDL and LOQ_blueberry_ were 0.412 and 1.092 μg/mL, respectively (Table 2).

**Table 2.** Method validation parameters for the quantification of monotropein.

| Parameter | Monotropein |
| --- | --- |
| Retention time | 1.04 min |
| Linear range | 0.04-40 µg/mL |
| Instrument detection limit (IDL) | 0.0103 µg/mL |
| LOQ<sub>LCMS</sub> | 0.0273 µg/mL |
| Method detection limit (MDL) | 0.412 µg/g |
| LOQ<sub>blueberry</sub> | 1.092 µg/g |
| Intraday recovery | 89-110 % |
| Intraday repeatability | 1.17-2.15 % RSD |
| Interday recovery | 91-108 % |
| Interday repeatability | 4.68-7.16 % RSD |

*Abbreviations: IDL = instrument detection limit; LOQ = limit of quantification; MDL = method detection limit; % RSD = percent relative standard deviation*

#### 3.2.2 Repeatability

Intraday and interday repeatability of monotropein are shown in Table 2. Intraday %RSD was 1.17-2.15%, while interday %RSD was 4.68-7.16%.

#### 3.2.3 Recovery

Recoveries of monotropein are shown in Table 2. Intraday recoveries were 89-110%, while interday recoveries were 91-108%.

### 3.3 Monotropein quantification in 28 blueberry species

The optimized method was applied to the quantification of monotropein in 28 blueberry fruit samples, including 15 wild and 13 cultivated varieties (Fig 1; Table 3). The range of monotropein content in wild *Vaccinium* species was 94-1807 ng monotropein/mg blueberry dry weight (dw), while in cultivated varieties monotropein ranged from below detection limits to 1460 ng/mg dw. Among the varieties analyzed here, wild *Vaccinium* species contained a significantly higher average concentration of monotropein than cultivated varieties (813 vs. 364 ng/mg dw; *p* = 0.0006), though several cultivated varieties contained more monotropein than some wild species.

**Fig. 1.**
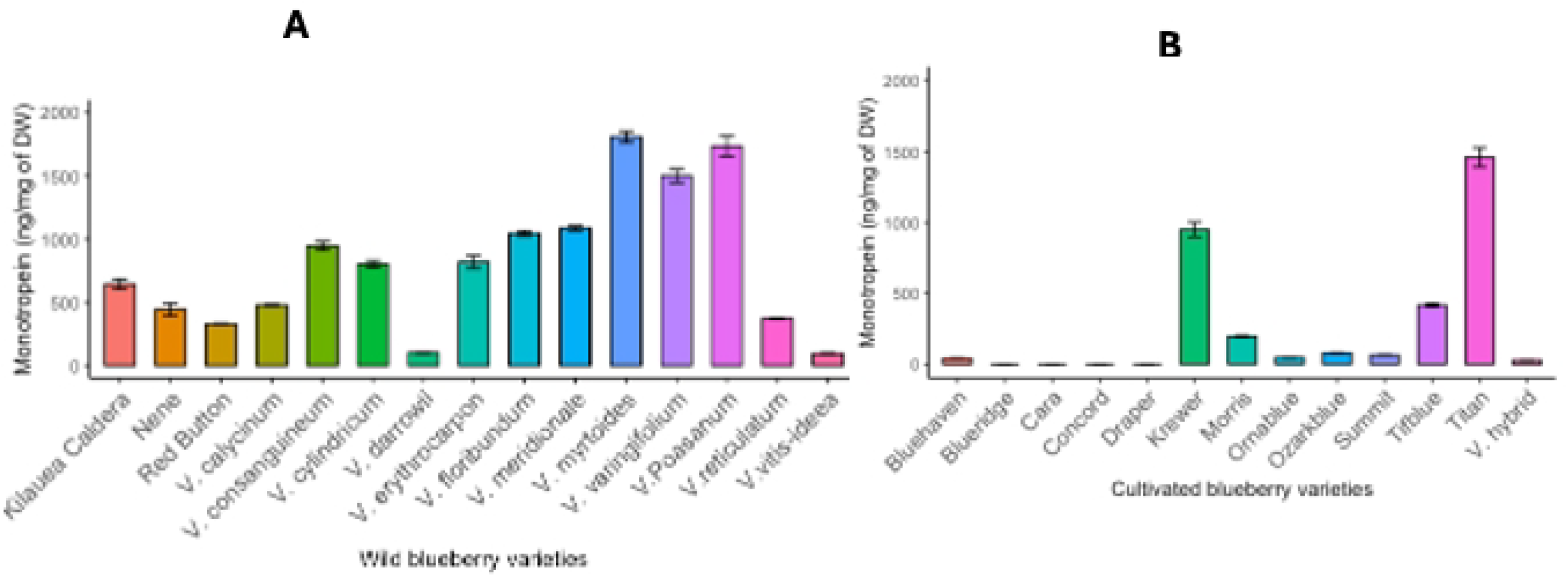
Monotropein content (μg/mg DW) in blueberries. A) Wild *Vaccinium* species and B) cultivatd blueberries. Error bars represent standard deviation (SD) from 3 technical replicates.

**Table 3.**
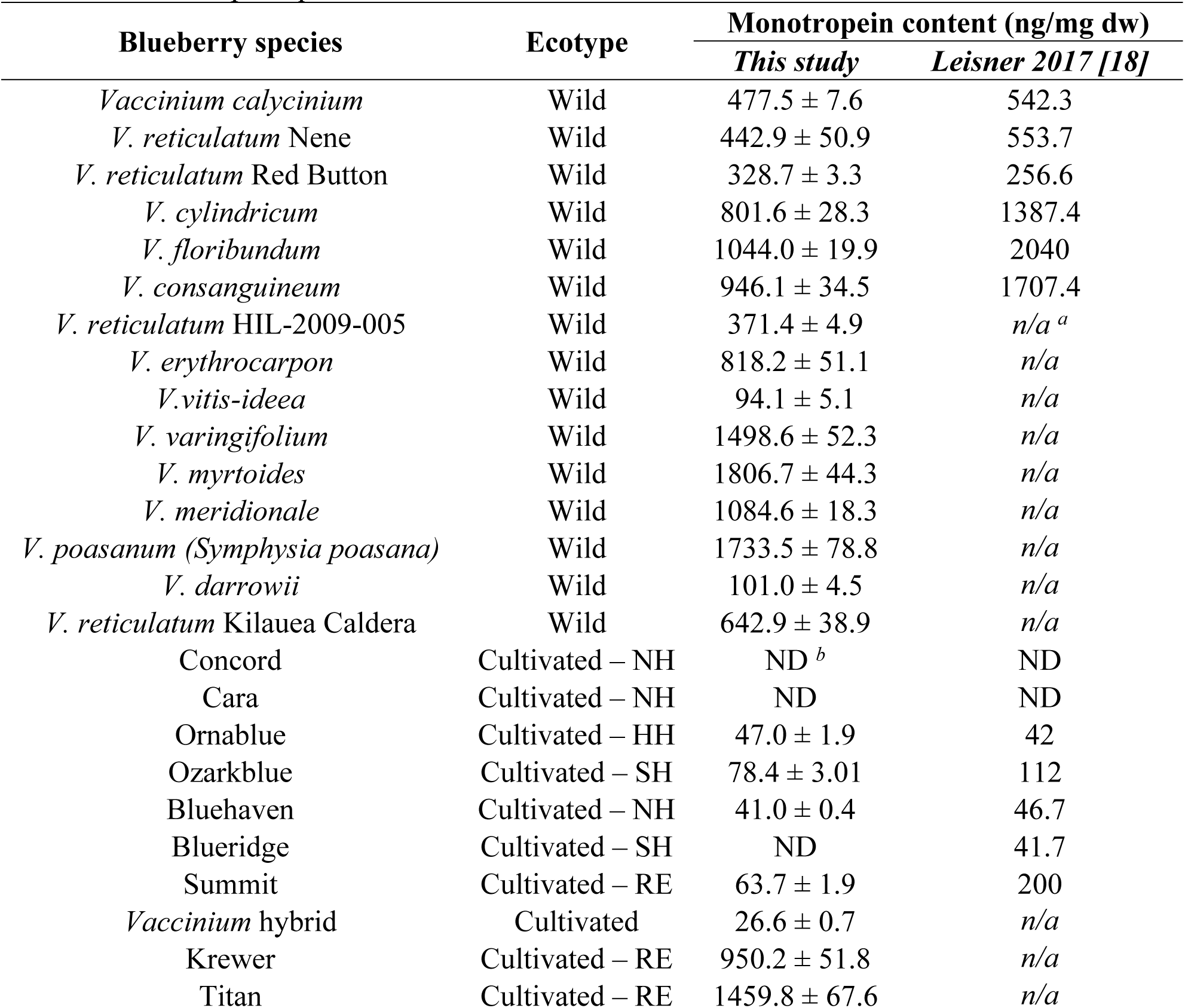
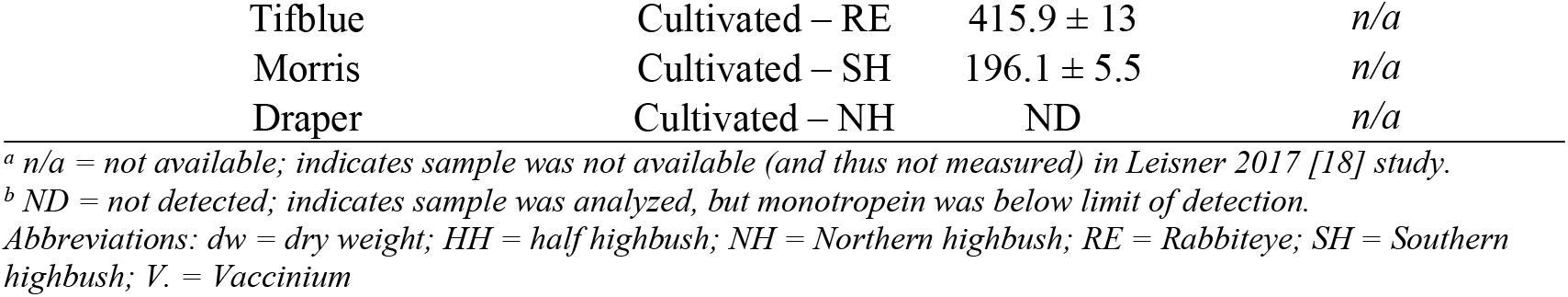
Monotropein quantification in blueberries.

## 4. DISCUSSION

In this experiment, the extraction and quantification of monotropein in lyophilized blueberry fruits was optimized and applied to the analysis of 28 blueberry cultivars (15 wild and 13 cultivated varieties). The most efficient extraction method was determined to be heating at 60°C for 15 minutes. When evaluating longer extraction times and the potential use of sonication (with or without heat), there were no significant differences in extraction efficiency, though lower temperatures did decrease extraction efficiency. The final conditions are shorter than those previously reported [18] while also improving on the previous method to provide quantitative recovery of monotropein from blueberry fruits, as demonstrated by the high intra- and interday recovery values. The method appeared to have good sensitivity, with IDL 0.0103 µg/mL, MDL 0.412 µg/mL, LOQ_LCMS_ 0.0273 µg/mL, and LOQ_blueberry_ 1.092 μg/mL, resulting in the quantification of monotropein in 24 of the 28 samples evaluated.

To contextualize the results of the present study, they were compared to the results of monotropein in cultivated and wild blueberry fruits from Leisner et al [18]. Both investigations showed that wild species had more monotropein, on average, than cultivated varieties. However, while the range of monotropein in wild *Vaccinium* species was similar between the two studies (94-1807 ng/mg dw (present study) vs 20-2371 ng/mg dw [18]), the results for cultivated species were different. The Leisner et al. investigation reported that monotropein in 66 of 71 cultivated varieties (i.e., 93%) was below the limit of detection, and of the 5 cultivated varieties that did contain quantifiable monotropein, the maximum observed value was 180. ng/mg dw [18]. In the present study, monotropein was below the limit of detection in only 4 of 13 cultivated varieties (i.e., 31%), and the 9 cultivars containing quantifiable monotropein averaged 364 ng/mg dw, which is double the amount reported in the highest cultivated variety in [18]. This is likely due to differences in the ecotypes analyzed in both studies, as the cultivars in [18] were primarily Northern Highbush (NH). As previously reported, cultivated varieties with wild *Vaccinium* parentage are more likely to contain monotropein [18]. Differences in the wild *Vaccinium* parentage in NH compared to southern highbush (SH) or rabbiteye (RE) ecotypes may explain why non-NH ecotypes are more likely to contain quantifiable levels of monotropein [18]. In the present study, the cultivated blueberries consisted of 4 NH and 9 non-NH ecotypes. Monotropein was only quantified in 1 of 4 NH cultivars, while it was quantified in 8 of 9 species belonging to other ecotypes, which aligns well with the hypothesis that the presence of monotropein in cultivated varieties is due to introgressions from wild species.

Interestingly, when comparing the 13 varieties (6 wild and 7 cultivated) that were analyzed in both [18] and the present investigation, monotropein levels were similar or slightly lower in the current samples (Table 3). The similarity across both studies provides additional support for the consistency of the method across multiple lab settings. Where differences were noted, there are likely due to environmental factors, including harvest year, soil composition, and both micro- and macro-environmental conditions, as prior studies have highlighted the role of environmental factors in modulating the production of specialized metabolites in plants [1, 29-31]. Other potential sources of variation include differences in seed source, sampling time, and interlab variations.

### Future Directions

Future research should prioritize a deeper investigation into the influence of gene– environment (G×E) interactions on monotropein biosynthesis and accumulation in blueberry. Teasing out the relative contributions of genetic background, environmental factors (such as soil composition, climate variability, and cultivation practices), and their synergistic effects will be critical to understanding the observed variability in monotropein levels among cultivars and wild accessions. While most studies on monotropein biosynthesis have been conducted in *Morinda* species, there remains a significant knowledge gap regarding the regulatory mechanisms and metabolic pathways governing monotropein production in *Vaccinium* spp. In *Morinda*, monotropein concentrations can be several-fold higher [14], highlighting the need to explore whether similar biosynthetic potential exists in blueberry through targeted breeding or biotechnological approaches. Additionally, future efforts should focus on evaluating the metabolic stability, transformation, and bioavailability of monotropein in the human body following blueberry consumption. Such studies will help clarify the extent to which monotropein contributes to the health-promoting properties of blueberries. Ultimately, integrating metabolomic profiling with genomics and breeding strategies will enable the development of cultivars with optimized monotropein content, thereby enhancing their functional and commercial value in the nutraceutical market.

## 5. CONCLUSION

In this study, methods to extract and quantify monotropein in blueberries was successfully developed, validated, and applied to the analysis of 28 blueberry cultivars, 15 of which were analyzed for the first time. The results of the present study aligned well with those of a previous investigation of blueberry monotropein, supporting the observations that wild *Vaccinium* species contain higher levels of monotropein than cultivated varieties and that cultivated ecotypes with introgressions of wild *Vaccinium* species are more likely to contain monotropein. As monotropein has been linked to potential health benefits, understanding the biosynthetic pathway to produce monotropein may aid breeding programs in enhancing the monotropein content of blueberries. Future studies should focus on the understanding the molecular mechanisms that produce monotropein to increase the nutritive value of blueberries.

## ACKNOWLEDGMENTS

This work is supported by the Foundational Knowledge of Plant Products program, project award no. 2022-67013-36416, from the U.S. Department of Agriculture’s National Institute of Food and Agriculture. This work was also supported by UDSA-NIFA Hatch project VA-160221.

## CONFLICT OF INTEREST

The authors state no conflict of interest with this work.

## AUTHOR CONTRIBUTION

Conceptualization - IK, CPL and DPC; Methodology-IK and DPC; Validation – IK, DPC; Formal Analysis-IK; Investigation-DPC, CPL; Writing-IK, CPL and DPC; Review & Editing-IK, CPL and DPC; Visualization – IK, DPC and CPL; Funding Acquisition-CPL and DPC.

## SUPPLEMENTAL INFORMATION

**Table S1**. Information of the accessions used in the study.

**Supplemental Figure S1. Sample multiple reaction monitoring (MRM) chromatograms for monotropein using an authentic standard**. Four transitions were measured; 413.117 → 233.036 was used for quantification.

**Supplemental Figure 2. Sample multiple reaction monitoring (MRM) chromatograms of monotropein in wild blueberry composite** (WBC). Values represent extraction using wild blueberry powder obtained from WBANA. A total of four mass transitions were optimized using IntelliStart software (413.117 → 185.020, 413.117 → 202.978, 413.117 → 233.036, and 413.117 → 251.056.

## Notes

### Competing Interest Statement

The authors have declared no competing interest.

